# Clean Air Travel: Evaluation of Mask-Free Alternatives to N95 During the Delhi “Air Emergency”

**DOI:** 10.64898/2025.12.05.692625

**Authors:** Devabhaktuni Srikrishna

## Abstract

**Background:** Air travelers, healthcare workers aiming to prevent infection and pollution-sufferers (e.g. Delhi “Air Emergency”) may experience difficulty during prolonged use of respirators (e.g. N95) incompatible with eating/drinking/sleeping. Protection factor of 5x-1000x (80%-99.9%) is targeted for infection prevention. Daily PM 2.5 time-series data suggest at least 4x-5x (75%-80%) protection factor 24×7 during the worst stagnant pollution days may reduce daily inhaled exposure to annual averages in Delhi.

**Methods:** Handheld Do-It-Yourself test methods were developed using a 7-channel optical particle counter. Three baseball cap visor-mounted air purifiers were prototyped enabling eating/drinking, “Air” (1 lb), “Pro” (2 lb), “Max” (2 lb 10 oz), and one bed-mounted prototype, “Sleep”. Clean air delivery rate per watt was measured for vertically stackable portable air purifiers to optimize power/floorspace/cost-efficiency.

**Results:** At 0.3 µm at nose-level, “Air” measured 87%, “Pro” measured 94%, “Max” measured 98%, a 2x-10x improvement in particle penetration over prior wearable purifiers (unobstructed in horizontal and lower visual fields) reporting approximately 60% (< 70%) of 0.3 µm particles at nose level. At 1.0 µm and 5.0 µm at nose-level, they measured 94–99% and 95–99% respectively. “Sleep” measured 91% at 0.3 µm at nose level. Stackable air purifiers measured 6-19 ACH in ∼3000 cubic foot room (300-900 cubic feet per minute) at 20-220 watts, with highest power-efficiency 19 CFM/Watt at 962 CFM.

**Conclusions:** In principle, maskless alternatives to N95 combine with floorspace- and power-efficient stackable/sleeping air purifiers for protection factor of 99%-99.98% (100x-5000x) in airplanes, submarines, ships, trains, hospitals, shelters with limited floorspace (e.g. berths).

## Introduction

### Difficulties of use of masks/respirators for infection prevention during travel and prolonged use

Sports medicine physicians reported 5–10% of athletes could contract viral acute respiratory infections during wintertime air travel [1] [2]. In 2020, it was observed that despite high-efficiency filtering inflight, passengers nearby an index case are at higher risk of infection of SARS-CoV-2 [3]. A systematic review of aircraft-acquired SARS-CoV-2 transmission found passenger infection risk was flight duration-dependent, 25 times higher on long-haul flights versus shorter flights, and each hour increased the incidence rate by 1.53-fold [4]. N95 and KN95 are often referred to interchangeably, but across a variety of manufacturers in one study the average filtration efficiency of N95 respirators at 0.3 μm particle size was found to be much higher at 98% and more consistent than KN95 respirators at 81% [13] and California Department of Public Health noted that N95 with head straps provides better fit than KN95 or KF94 which use earloops [14]. Although KN95 (∼80% or 5x) to well-fitted N95 (∼95% or 20x) masks are used by some passengers inflight they are incompatible with eating/drinking and all-day/prolonged use for many people, and in 2020 the former CDC director Dr. Walensky commented about discomfort and suffocation with N95 masks, making them difficult to tolerate for long durations [5]. Lightweight respirators include N95/N99/FFP3, but difficulties wearing them (non-compliance) were also anticipated by Dr. Wiener in 1996 and the problem has remained unsolved [6]. In 2024, the JPEO-CBRND issued a request for information for “low-burden respiratory protection” for biological/pandemic threats [7]. During COVID-19 pandemic hearings, UK Health Security Agency testified FFP3 respirators were too uncomfortable for sustained use by healthcare workers [8] [9] and one study pointed to staff-to-staff transmission in tea-rooms (unmasking to eat/drink) as a possible hotspot of transmission during mask removal [10]. A draft CDC review concluded difficulty breathing, headaches, dizziness, skin barrier damage, itching, fatigue, and difficulty talking were more frequently reported among N95 respirator users [11]. Respirators for infection prevention also cannot be worn by patients at dentist due to need for direct access to the patient’s mouth. Even if masks are perfectly functional, their protection fails when removed for eating, heat stress, or discomfort during extended airborne contamination. Powered air purifying respirators (PAPR), especially those which are loose-fitting, are meant to overcome some of these challenges especially related to heat but often require a backpack and still facially obstructive, cumbersome, and/or noisy to use [12] and they are unsustainable for long-term, daily use otherwise they could be worn continuously like clothes or eyeglasses.

### Delhi, India: inconsistent respirator utilization during extreme, prolonged particulate pollution

Abruptly in November, 2025, Delhi has been described by some doctors as a “gas chamber” with patients streaming into hospitals with respiratory difficulties such as cough, asthma, and bronchitis [13] [14]. In one recent study as narrated by a doctor on national TV, of 4,000 children aged 11-14 years across 22 schools nearly 60% reported respiratory symptoms of persistent cough, sneezing, runny nose, chest pain, tightness, itching in eyes, etc. and one third with asthma [15]. Delhi, India during winter months is a case study of intense, prolonged, unpredictable cycles of air pollution, called an “Air Emergency” by local media. Patients are coughing due to pollution and doctors recommend N95/FFP2 when however they cannot be worn all the time due to a variety of discomforts and limitations [16], and a patient using a surgical mask reports although she has an air purifier she cannot use it outside the home (at 2:20 of [17]). A newspaper reported 80 per cent households in Delhi have had at least one member fall ill due to toxic air in the past month [18]. Even most AQI protestors were seen unmasked and others taking a break from masking with their masks below their chin, 10:45 of [19].

The primary concern is particulate matter pollution (PM 2.5) at 200-500 μg/m^3^ or more for days at a stretch with little relief in between. As seen on AQI.in^TM^, changes in particulate concentrations that that triggered the Delhi crisis were between ∼100 μg/m^3^ in late October to ∼300 μg/m^3^ in mid-November [20]. As seen on Purple Air^TM^, during Thanksgiving week in 2025, raw PM 2.5 in New Delhi within approximately 10x of San Francisco which is not reporting such a respiratory health crisis [21]. Such high levels of PM 2.5 are typically seen nearby wildfires (e.g. Los Angeles fires of 2025) which usually last for days until the trees burn up, but not recurring for months like in Delhi. Measurable increases in pulmonary illness, myocardial infarction requiring emergency medical attention were observed in the 90 day period following the LA fires [22].

### Protection factor targets for particulate pollution (5x-20x)

Anecdotal reports suggest November, 2025 in Delhi has been markedly worse than prior years [23] making an evening walk difficult [24] [25]. However, Daily PM 2.5 measurements over five years across 39 stations spread across Delhi metro area reported in Figure 4 of [26] show peak particulate levels roughly comparable to those observed this season and that if particle exposure is cut down in a sustained manner by a factor of:

- 4x-5x (75%-80%), even on most of the worst days particulate dose may be around the year-round average level of 100 µg/m^3^ is about four or five times lower than the peaks up at about 400-500 µg/m3 (Figure 3, Figure 4 of [26]).
- ∼10x (90%), which is the blue line shown in Figure 4 of [26] even on most of the worst days particulate dose may be around the national daily standard of 60 µg/m3,
- ∼20x (95%) which is the yellow line shown in Figure 4 of [26] even on most of the worst days particulate dose may be around the even more stringent WHO daily guideline of 15 µg/m3.

A study of particle size of pollution in Delhi reported most of the volume of particulate matter is between 0.1 to 1.0 micron (“Accumulation”) and 2.5 microns to 10 microns (“Coarse”) as shown in Figure 1 of [27], suggesting protection factors would at least need to apply to these size fractions [28] [29].

**Figure 1:**
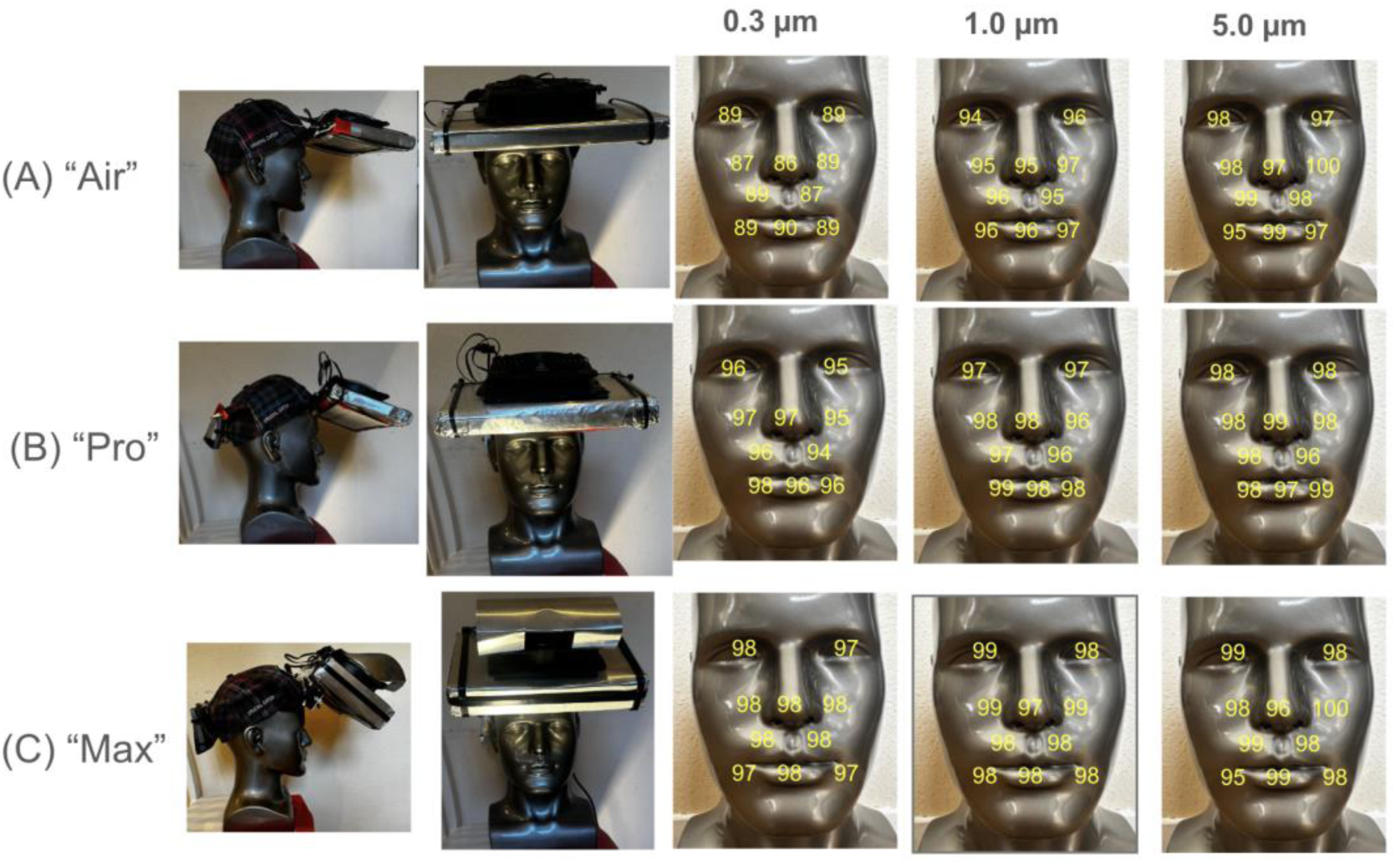
(A) “Air” using Arctic P12 and MERV 13 (6”x9”) with its pleats are oriented perpendicular to direction facing user, tilted using friction hinges attached to brim of hat (B) “Pro” similar to “Air” except with more powerful Arctic P12 Max fan and a second flattened MERV 13 filter below first pleated MERV 13 filter and counterweight in back (C) “Max” is similar to “Pro” except it uses two pleated 6”x9” filters and brackets to raise the assembly closer to the head, adds rain guard above fan

### Protection factor targets for infectious viruses/bacteria (5x-1000x)

In 1996, Dr. Wiener called for lightweight respirators for bioaerosols (excluding vapor protection) wearable for prolonged periods [30] due to lack of early warning of biological attacks, airborne contagion, and difficulty of sustaining gas-mask use beyond 3-4 hours specifying protection of factors of 1000x (99.9%) for aerosol particles above 1 µm and 50x-100x (98 to 99%) for particles below 1 µm. If present, infector’s quanta emission rate (1 quanta is minimum infectious dose inhaled) may range from 0 to over 1000 per hour [31] [32] requiring commensurate protection factors to prevent transmission, and ∼10 hour exposure (e.g. long flights or on ships/submarines at sea) further multiplies protection factor needed to prevent contagion by an order of magnitude [33] [34] [35]. The supplementary materials include infection risk modelling using the Wells-Reilly equation and well-mixed assuption for an infector/susceptible in enclosed spaces for different infector quanta emission rates, durations of exposure (1 hour versus 8 hours), clean airflow rate in the enclosed space, and with/without use of 95% effective masks.

However, at short distances in the “jet zone” from an infector [36] there is risk for pathogen transmission which likely exceeds the well-mixed assumption e.g., consider someone smoking a cigarette [37] [38]. Lack of ability to physically distance and prolonged duration of exposure likely contributes to transmission risk (e.g. in sleeping quarters in ships/subs/trains) [35]. In 2020, the US Navy reported that roughly 35% of sailors infected with COVID-19 among the uniformed navy population had “few to no symptoms” [39]. For example, in 2020, of approximately 5,000 sailors onboard the aircraft carrier USS Theodore Roosevelt, 416 tested positive for COVID-19 and 229 (55%) were reported asymptomatic [40], resulting in the ship returning to shore. A similar situation was described on a French aircraft carrier, Charles de Gaulle [33]. In 2020, SARS-CoV-2 spread aboard a nuclear submarine resulting in half the submariners eventually infected within 11 days in spite of the air aboard the submarine being continuously scrubbed [34]. Reflecting the increased risks of transmission from extended durations (e.g. 8 hours as modeled in Figure S2), in 2020 the US Navy recommended that sailors should sleep in alternating head-to-foot positions where berthing configuration allows [39], 14-day restriction of movement (ROM), testing, and “bubbles” without reporting another serious outbreak [41]. The Marine Forces Special Operations Command highlighted bubbles can still be breached by partner forces [42] or potentially by peers during littoral submarine operations. ROM precautions were subsequently relaxed for vaccinated sailors [43].

For infectious bioaerosols, there remains considerable debate and uncertainty about the relative importance of submicron versus micron particles. One reason is the percentage of infectious virus/bacteria in submicron versus micron particles (< 1.0 µm versus > 1.0 µm) depends on several competing and unknown factors that depend on pathogen, host, and environment (exhalation from deep lung versus upper respiratory pathway, evaporation or condensation before inhalation, temperature, humidity). For example, in one study, more of the viable virus from human subjects was observed in submicron respiratory aerosols than micron and above [44], whereas in another study most of the genetic material was captured in micron and above [45].

### Asymptomatic or Subclinical Contagion Risk

24×7 protection is needed to prevent asymptomatic, contagious viruses/bacteria which can infect without early warning from obvious symptoms (e.g. SARS-CoV-2) via lungs or eyes especially if prolonged or at close-range [46]. In a challenge trial inoculating healthy, unvaccinated young adults (ages 18 to 30 years) with SARS-CoV-2 and measuring their viral emissions using an air sampler, the extent of symptoms did not correlate well with measured viral emissions, and an asymptomatic individual was found to emit large amounts of virus for several days, including viable virus within a mask [47]. In a study documenting local transmission among Marine recruits during a supervised 2-week quarantine period, most who tested positive for SARS-CoV-2 were asymptomatic [48]. In 2021, 18% of 72 high school classrooms in South Africa had tuberculosis, likely subclinical, only discovered through bioaerosol sampling [49].

### Handheld Testing

For aerosols, difficulties with long-term use of respirators are often caused by facial obstruction and/or face seals that also obstruct eating and drinking. The 0.3 µm particle size is known as approximately the “most penetrating particle size”, whereby the aerosolized particle contamination filtration efficiency is typically lower than substantially larger particles [50] for which calibrated, handheld optical particle counters are readily available. Subsequently testing of N95 masks by NIOSH has been updated to use a size of about 0.075 µm [51], but regardless of which size is used if the mass fractions of particulates below 0.3 µm are expected to be minimal or their filtration efficiency is expected to be higher, testing at 0.3 µm in principle provides reasonable assurance of particle filtration.

In other publications wearable aerosol filtration devices either above or below the breathing zone, some have reported 95% or greater for particles sized around 0.3 µm directly at the output of the air filtration but have apparently not reported 95% for particles sized around 0.3 µm at nose level from submicron contaminants entering the breathing zone without occlusion/obstruction of the horizontal and lower visual field needed to maintain forward and ground-visibility [52] [53] [54] [55] [56] [57].

In shelters or enclosed spaces such as an airframe, steady-state protection factor is ratio of air filtration rate to contaminated infiltration rate into the room (expressed in cubic feet per minute or CFM, cubic meters per hour, or air changes per hour, ACH), with infiltration dynamically dependent on sealing of leaks, outside wind, and opening/closing windows and doors. For example [35] uses the same handheld test as used in this research to measure ACH:

- 9 ACH of aerosol filtration, within 40 minutes after closing the windows resulted in 50-fold (98%) reduction (protection factor) observed in a test room for particles having a size of 0.3 µm relative to the outdoor concentration
- ACH on three passenger jets inflight (Airbus A319, Airbus A321, Boeing 737 Max 8/ 9) were measured the range of 11 to 12 ACH.

Imperfectly sealed shelters can experience high infiltration rates (0.1-1.0 ACH or more), requiring over 10 ACH of filtration for 90–99% exposure reduction. A higher air changes per hour (ACH) rate in a room increases the protection factor against aerosol spikes by reducing contaminant concentrations more rapidly, thereby lowering the dose inhaled by occupants. One exception is at short distances in the “jet zone” from an infector [36] there is risk for pathogen transmission e.g., consider someone smoking a cigarette [37] [38].

## Methods

### Maskless Protection Devices: Construction, Materials, Manufacturing, and Supply Chain

This study did not involve participation by human subjects, and IRB approval is not needed. We investigated, developed, and evaluated the protection factor of maskless prototypes: facially-unobstructed respirators designated “Air,” “Pro,” “Max,” as mounted on a standard mannequin head shown in Figure 1 (A) to (C) based on US Patents # 12,343,573-B1, 12,496,474-B1. They were assembled and tested using cap visor-mounted, lightweight fans drawing externally contaminated air through filters, balancing filtration efficiency with air resistance to modulate the aerosol field around breathing and ocular zones. They permit eating/drinking without occlusion/obstruction of the horizontal and lower visual field needed to maintain forward and ground-visibility. The devices were cut, assembled, and tested using mass-manufactured materials (e.g. aluminum sheet), PC case fans (12-volt DC), and HVAC filters from commercial-off-the-shelf (COTS) supply chain, not dependent on custom manufacturing techniques (e.g. injection molding), and manufactured for large secondary markets with established demand from consumers and businesses for supply chain resilience in case of a demand-surge as seen for N95 during 2020 when they became unavailable. Although prototypes were hand-assembled, they are not dependent on specialized manufacturing techniques (e.g. injection molding) and can transition to assembly-line production based on availability of COTS components.

Figure 1 (D) shows a prototype “Sleep” also based on US Patents # 12,343,573-B1, 12,496,474-B1 which is a sleeping respiratory protection device to provide clean air to the breathing zone while asleep. The figures show the frontal and rear views of the device with a standard mannequin, based on an air processing assembly that consists of two 24”x12” pleated MERV 13 filters arranged in series (1” thick 3M^TM^ MERV 13 and 4” thick Nordic Pure^TM^ MERV 13), ten Arctic^TM^ P12^TM^ fans arranged in parallel on a 5×2 grid, a fan grille, and canopy to guide clean airflow toward and over the face of the user and exclude contaminated air. The fans were powered by a 12V AC/DC adapter.

For aerosol filtration, all cap visor-mounted devices relied on aerosol filters cut from approximately 0.75” thick pleated HVAC filters rated MERV 13 or MERV 14, either one or two in series made by 3M^TM^ (Filtrete^TM^) cut to 6”x9”. The additional flat filter in “Pro” was flattened from a pleated MERV 13 filter made by Lennox^TM^.

The fans used in the maskless protection devices were either P12^TM^ or P12 Max^TM^ made by Arctic^TMs^ and powered by 12V DC portable batteries.

### Supplementary Air Filtered Exchanges (SAFE): Construction, Materials, Manufacturing, and Supply Chain

For an indoor air filtration system, efficiently achieving a target air changes per hour (ACH) or filtration rate requires scaling up the indoor air filtration system’s clean air delivery rate (CADR) in units of cubic feet per minute (CFM) without excessively scaling up any of the trifecta of operational resource burdens, i.e., the power consumed in watts, noise generated in dBA, and floorspace / volume taken up in square feet and cubic feet respectively. Conversely, an indoor air filtration system becomes unscalable/unsuitable especially for larger rooms if designed inefficiently in terms of the following metrics: CFM per watt, CFM per dBA, CFM per square foot, or CFM per cubic foot.

To overcome these resource consumption challenges for indoor air filtration systems and reach the protection factors needed for control of infectious bioaerosols (multiplicatively with maskless protection devices), we also developed portable air purifiers Supplemental Air Filtered Exchanges (**SAFE-1**), quadruply stacked, easy-to-assemble from pleated MERV 13-16 HVAC filters and Lasko^TM^ fans previously described as SAFE in [58] from commercial-off-the-shelf (COTS) supply chain and tested singly in [59]. SAFE-1 on lowest speed is approximately 55 watts per fan (120-volt AC power), is vertically stackable to make most efficient use of floorspace, and wall-adjacent, wall-mountable power outlets access near the walls of a shelter. In single units, SAFE have been in daily use in at least four schools and several homes in San Francisco since 2022 and clean air delivery rate verified by a researcher at CDC [60]. MERV 13-16 filters remained effective during daily usage during the school days in an elementary school for an academic year [35] [61]. To improve on SAFE-1, prototype SAFE-2 was developed attaching cylindrical shrouds to SAFE-1, and SAFE-3/4 with PC fans (12-volt AC), box fans (120-volt AC) all of which are stackable to multiply their airflow rate.

- **SAFE-1:** Figure 2 (B) shows four vertically stacked DIY air filters as described in [58]
- **SAFE-2:** Figure 2 (C) shows the same setup as SAFE-1 but with a 12” cylindrical shroud attached to the output of each fan to enhance air mixing.
- **SAFE-3:** Figure 2 (D) shows ten Arctic^TM^ P12^TM^ case fans (12 volts DC) in a grid formation and two Nordic Pure^TM^ MERV 13 filters (4”x12”x24”)
- **SAFE-4:** Figure 2 (E) shows fifteen Arctic^TM^ P12^TM^ fans in a grid formation (5×3) and two Lennox^TM^ MERV 13 filters (5”x16”x24”)
- **SAFE-5:** Similar to SAFE-4, except with Arctic^TM^ P12^TM^ Max fans

**Figure 2:**
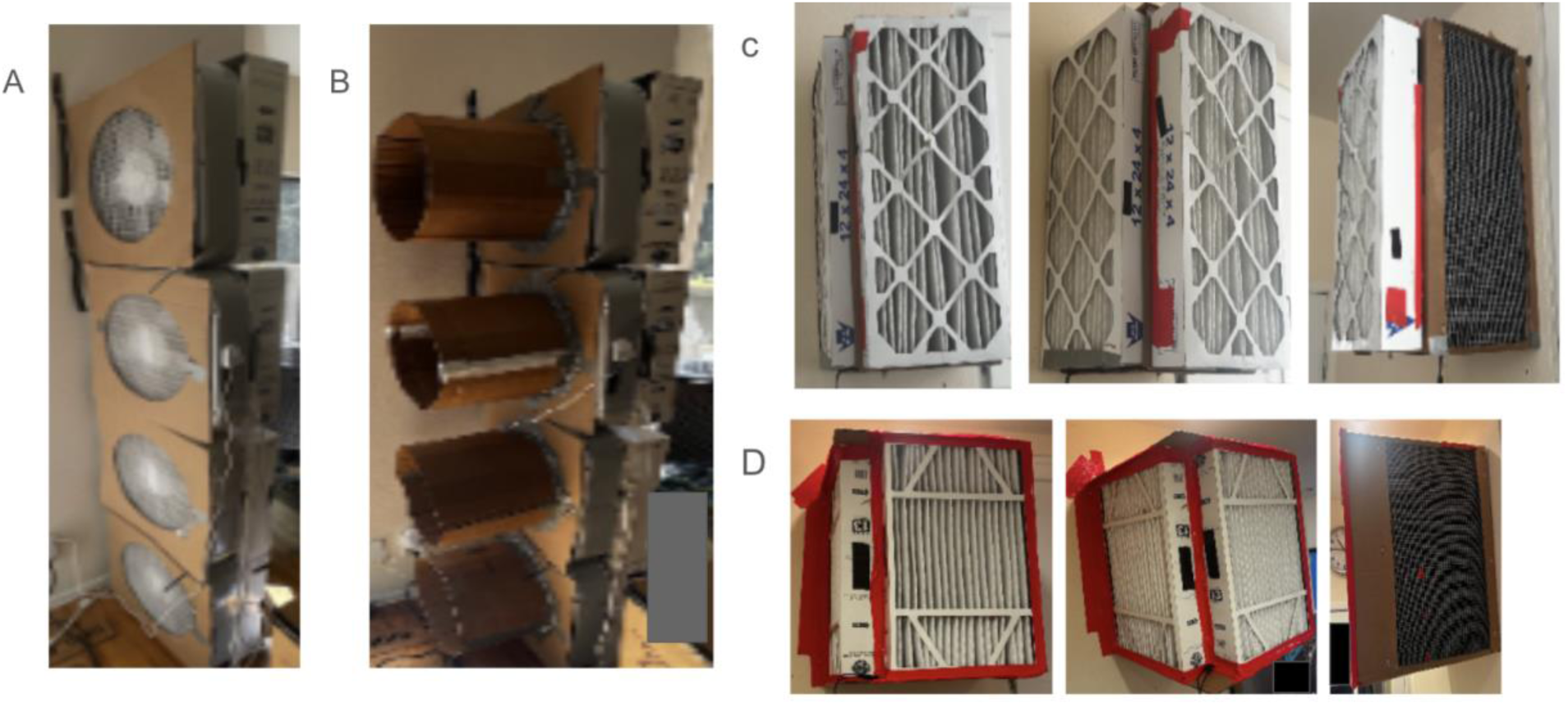
(A) SAFE-1 (B) SAFE-2 (C) SAFE-3 (D) SAFE-4.

### Instruments

Instruments used to take measurements in the experiments are listed below:

- OPC1: Particle counts were measured with generated smoke aerosols (and ambient aerosols) using a handheld optical particle counter calibrated by the manufacturer (Temtop^TM^ PMD 331^TM^) which reports particle counts per liter across 7 channels: 0.3 µm, 0.5 µm, 0.7 µm, 1.0 µm, 2.5 µm, 5 µm, and 10 µm [59]. A hose was used to precisely direct the particles from each location on the face to the input of the particle counter as shown in Figure 3.
- OPC2: In some instances, as secondary confirmation, particle counts at 0.3-0.5 µm were verified using a second optical particle counter (TSI^TM^ Omnitrak^TM^ PM module) which reports particle counts per liter across five channels: 0.3-0.5 μm. 0.5-1.0 μm, 1.0-2.5 μm, 2.5-4.0 μm, 4.0-10 μm. Since the input to this particle counter is large (2”x1”), use of a hose was not feasible.
- FTM: Airspeed was recorded using a vane anemometer (BTMETER^TM^ BT-100^TM^ Handheld Anemometer) measured in feet per minute (FTM).
- DBA: Noise was measured using the NIOSH Sound Level Meter app on an iPhone 14 in decibels above audible (dBA).
- OZ: Weight was measured using a standard desktop weight scale (Escali^TM^) in ounces.

**Figure 3:**
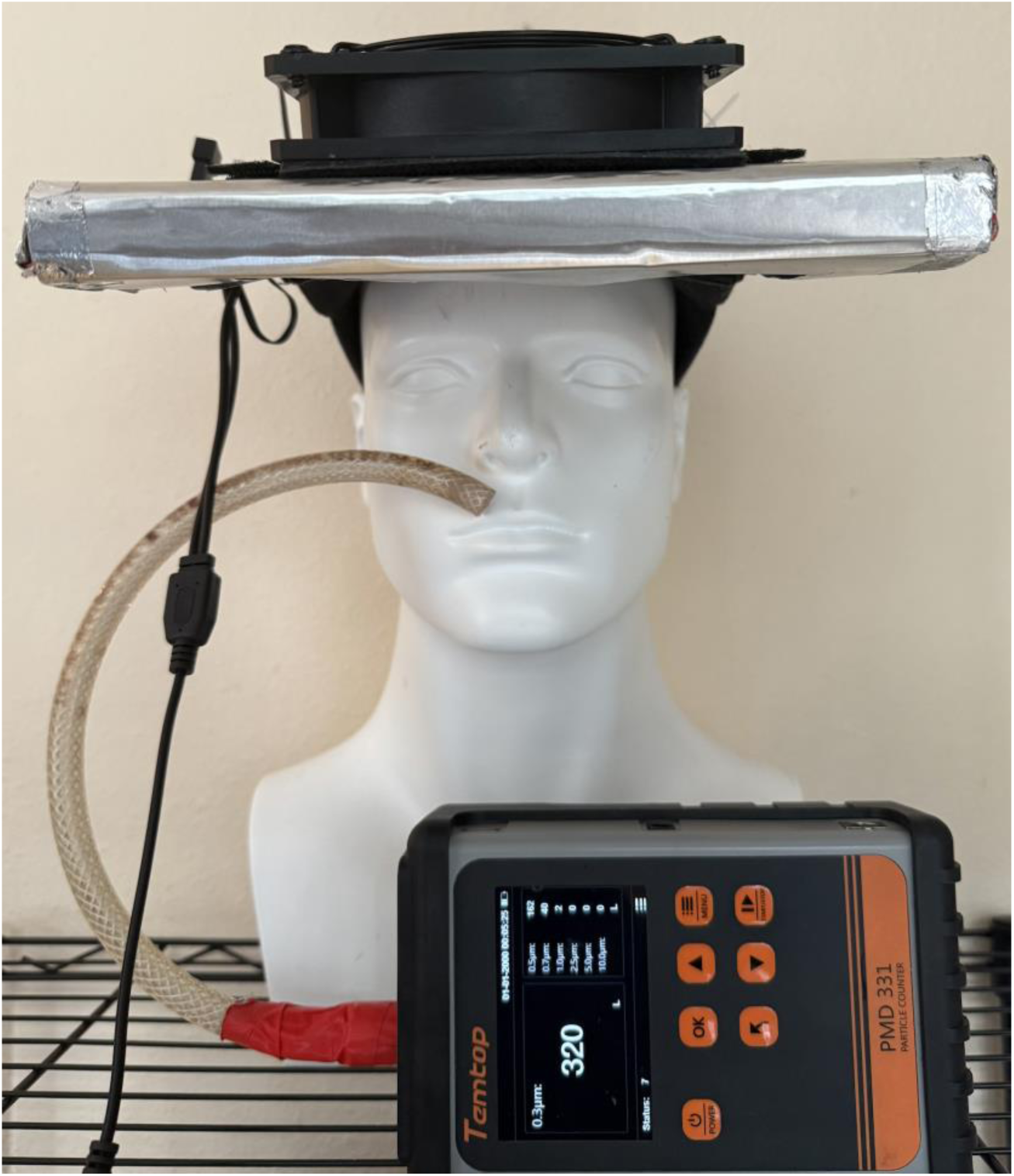
OPC1 with intake input of hose positioned between the right nostril and mouth of mannequin head.

All tests were conducted indoors at room temperature of approximately 74 degrees Fahrenheit.

### Handheld Test Procedure for Maskless Protection Devices

Filtration efficiency for aerosol concentrations was computed using the formula, 1 - location concentration / intake concentration:

- OPC1 was used to compute the filtration efficiency at 0.3 µm at each of the 10 locations on the face of the mannequin by recording for 10 seconds sequentially. Intake particle concentration (particles per liter) above the device (near the fan intake) were recorded also for 10 seconds, before and after recording at each of the 10 locations and then averaged.
- For secondary confirmation, OPC2 was used to compute the filtration efficiency at 0.3-0.5 µm, for 5 minutes at nose level while the fan was on, and before and after when fan was off also for a duration of 5 minutes each and averaged.

Filtration efficiencies with OPC1 were measured five times and averaged, except for “Max” they were measured 10 times and then averaged. Airspeed at chin level and noise within 3 inches of the ear were measured five times and averaged. Figure 3 shows an example of the “Air” device with input of the hose connected to OPC1 positioned between the right nostril and mouth of mannequin head.

### Handheld Test Procedure for Air Changes Per Hour (ACH), noise, and Power, and Airspeed

Details of test procedure to measure ACH is similar to that published in Supplemental Appendix of [35] and also included in Supplementary Materials. Noise was measured for each test using DBA at a 9-inch distance from the fan perpendicular to the direction of the output airflow. Power consumed during each test was measured using WATT. Airspeed was measured at the output of the fan using FTM.

## Safety Notes

The resources and information in this article are for informational purposes only and should not be construed as professional advice. The content is intended to complement, not substitute, the advice of your doctor. You should seek independent professional advice from a person who is licensed and/or qualified in the applicable area. No action should be taken based upon any information contained in this article. Use of the article is at your own risk. Author (and Patient Knowhow, Inc.) are not associated with any of the manufacturers mentioned in this research, and assume no liability for any content made available in this article. Patented (US 12,343,573-B1, 12,496,474-B1) and worldwide patent-pending. All rights reserved.

## Results

### Test Results for Maskless Protection Devices

Details of the devices and measurements (and their standard deviations in case of multiple measurements) are in Table 1. Filtration efficiency for particles sized 0.3 µm, 1.0 µm, and 5.0 µm at 10 different locations across the face (nose, mouth, eyes) are shown for “Air,” “Pro,” and “Max” are also shown in Figure 1. A 20-second YouTube short [62] demonstrates a nebulizer to visualize the flow of water-based aerosols (representing external aerosolized contamination) at different points of the breathing zone when “Max” is turned on and off. First the nebulizer is positioned to introduce aerosols from below the nose, then it is introduced from in front of the nose. In both cases the aerosol can be seen to be deflected from the nose, mouth and eyes when the device is turned on.

**Table 1:** Measurements of maskless protection devices.

|  | ("Air")<br>Single MERV 13, 6"<br>x 9" | ("Pro")<br>Dual MERV 13<br>pleated/flat, 6" x<br>9" | ("Max")<br>Dual MERV 14, 6"<br>x 9" | ("Sleep")<br>Dual MERV 13,<br>24" x 12" |
| --- | --- | --- | --- | --- |
| Number of tests | 5 | 5 | 10 | 5 |
| Below left nostril (0.3 µm, 1.0 µm, 5.0 µm) | 87%, 95%, 98% | 94%, 96%, 96% | 98%, 99%, 98% | 91%, 92%, 88% |
| STDEV Below left nostril | 3%, 2%, 3% | 3%, 2%, 2% | 1%, 1%, 3% | 1%, 4%, 18% |
| Below right nostril (0.3 µm, 1.0 µm, 5.0 µm) | 89%, 96%, 99% | 96%, 97%, 98% | 98%, 98%, 99% |  |
| STDEV Below right nostril | 2%, 1%, 1% | 2%, 2%, 3% | 1%, 1%, 2% |  |
| Side of left nostril (0.3 µm, 1.0 µm, 5.0 µm) | 89%, 97%, 100% | 95%, 96%, 98% | 98%, 99%, 100% |  |
| STDEV Side of left nostril | 2%, 2%, 0% | 3%, 2%, 2% | 0.4%, 1%, 2% |  |
| Side of right nostril (0.3 µm, 1.0 µm, 5.0 µm) | 87%, 95%, 98% | 97%, 98%, 98% | 98%, 99%, 98% |  |
| STDEV Side of right nostril | 3%, 2%, 3% | 1%, 1%, 2% | 1%, 1%, 5% |  |
| Front of nose (center) (0.3 µm, 1.0 µm, 5.0 µm) | 86%, 95%, 97% | 97%, 98%, 99% | 98%, 97%, 96% |  |
| STDEV Front of nose (center) | 2%, 2%, 3% | 1%, 2%, 2% | 1%, 2%, 5% |  |
| Front of mouth (center) (0.3 µm, 1.0 µm, 5.0 µm) | 90%, 96%, 99% | 96%, 98%, 97% | 98%, 98%, 99% |  |
| STDEV Front of mouth (center) | 1%, 1%, 3% | 2%, 1%, 1% | 1%, 1%, 2% |  |
| Left of mouth (0.3 µm, 1.0 µm, 5.0 µm) | 89%, 95%, 99% | 96%, 98%, 99% | 97%, 98%, 98% |  |
| STDEV Left of mouth | 2%, 3%, 3% | 3%, 2%, 1% | 1%, 1%, 4% |  |
| Right of mouth (0.3 µm, 1.0 µm, 5.0 µm) | 89%, 96%, 95% | 98%, 99%, 98% | 97%, 98%, 95% |  |
| STDEV Right of mouth | 2%, 2%, 5% | 1%, 1%, 2% | 1%, 1%, 6% |  |
| Left eye (0.3 µm, 1.0 µm, 5.0 µm) | 89%, 96%, 97% | 95%, 97%, 98% | 97%, 98%, 98% |  |
| STDEV Left eye | 3%, 1%, 3% | 3%, 3%, 3% | 1%, 1%, 3% |  |
| Right eye (0.3 µm, 1.0 µm, 5.0 µm) | 89%, 94%, 98% | 96%, 97%, 98% | 98%, 99%, 99% |  |
| STDEV Right eye | 2%, 2%, 3% | 3%, 3%, 2% | 1%, 1%, 2% |  |
| Omnitrak PM (0.3-0.5 µm) | 90% | 98% | 97% |  |
| Airspeed (feet per minute) | 118 | 78 | 181 | 177 |
| STDEV Airspeed | 0 | 0 | 8 | 0 |
| Noise (dBA) | 50 | 62 | 62 | 55 |
| STDEV Noise | 1.5 | 0.7 | 0.7 | 0.2 |
| Weight (oz) | 15.6 | 32.0 | 41.8 |  |

**Table 2:** Measurements of SAFE.

| Example (length x width x height) | Fan Speed Setting | Aerosol CADR (estimated from ACH measured, number of data points, average percent error) | Vapor CADR (estimated from ACH measured, number of data points, average percent error) | Airspeed at Fan (feet per minute) | Noise (dBA) | Power (watts) |
| --- | --- | --- | --- | --- | --- | --- |
| SAFE-1: 4 Lasko™ 20" fans x two MERV 16 and two MERV 13 (9"x20"x82") | 1 | 869 CFM (17.6/hour, 21, 4.8%) | n/a | 511 | 60 | 220 |
| SAFE-2: 4 Lasko™ 20" fans x two MERV 16 and two Lennox™ MERV 13 (21"x20"x82") | 1 | 991 CFM (20.1/hour, 21, 7.2%) | n/a | -- | 60 | 220 |
| SAFE-3: 10 Arctic™ P12 fans x two 12"x24"x4" Nordic Pure MERV 13 (16"x16"x24") | n/a (maximum) | 330 CFM (6.70/hour, 21, 3.0%) | n/a | -- | 58 | 20 |
| SAFE-4: 15 Arctic™ P12 fans x two 20"x24"x5" Lennox™ MERV 13 (25"x25"x24") | n/a (maximum) | 592 CFM (12.0/hour, 21, 4.2%) | n/a | -- | -- | 30 |
| SAFE-5: 15 Arctic™ P12 Max fans x two 20"x24"x5" Lennox™ MERV 13 (25"x25"x24") | n/a (maximum) | 962 CFM (19.5/hour, 21, 10.8%) | n/a | -- | -- | 50 |

### Test Results for SAFE

A series of tests were conducted using SAFE-1 to SAFE-5. The test results are shown in Table 1.

## Discussion

Prototype maskless protection devices measured 87–98% aerosol filtration at 0.3 µm at nose level (2–10% exposure), which appears to be a 10x improvement over published wearable devices that do not occlude/obstruct the horizontal and/or lower visual field needed to maintain forward and ground-visibility and enable eating/drinking:

- Max with larger 6” x 9” MERV 14 pleated filters (two in series) was able to exceed 97% across all 10 locations and 98% in some locations representing the best, most uniform result for aerosol filtration efficiency across all the three devices tested.
- Pro, considerably lighter than Max, demonstrated 94%-97% across all 10 locations
- Air measured 50 dBA near the ear with some ambient noise resulting in filtration efficiencies more than 85% uniformly across all 10 locations. In contrast to 0.3 µm, Air measured 95% and 98% at 1.0 µm and 5.0 µm, respectively. “Air” (> 5x or > 80% at 0.3 µm) might be sufficient to reduce particulate exposure for air pollution.
- Sleep measured 55 dBA near the ear resulting in filtration efficiencies more than 85% at at 0.3 µm, 1.0 µm and 5.0 µm.

SAFE, Simple-to-assemble and PACs (vertically stackable) for aerosols-only measured 6-19 ACH (300-900 CFM aerosols) at 20-220 watts, with highest power-efficiency 19 CFM/Watt at 962 CFM.

- SAFE-1 and SAFE-2 using box fans demonstrate ACH of 17-20 (869-991 CFM) for a power efficiency of 4 to 4.5 CFM/Watt, also demonstrating how stacking can be used to scale up ACH.
- SAFE-3 and SAFE-4 using PC case fans (Arctic^TM^ P12^TM^) demonstrate ACH of 6.7-12 (330-592 CFM) for a power efficiency of 16.5-19.9 CFM/Watt, approximately four times the power efficiency than SAFE-1 and SAFE 2.

### Limitations

Theoretically, results from the test aerosols used in this study in some instances might not extrapolate directly to removal of aerosols with specific viruses/bacteria, different kinds of pollution, and particle sizes other than those at reported herein for 0.3 µm, 1.0 µm, and 5.0 µm (such as 0.075 µm).

Although this study included tests of aerosol filtration efficiency at 10 locations on the face of a mannequin, it did not attempt to measure the (in)sensitivity to varying head sizes, facial shapes, heavy breathing, eating, or drinking. This study also does not measure the effect of movement (e.g. walking, running), ambient wind currents, and strong air jets generated by exhalations from nose or mouth of another person facing the wearer’s face. Although the higher the air speed that the cap visor-mounted wearable air purifier generates at nose level the less affected it is likely to be.

One question is by how much the vertical airflow in cap visor-mounted wearable air purifiers provide protection from (or to) others who may be directly in front of the wearer during close-contact (e.g. < 3 feet) if the wearer’s breathing and exhaled particles (or those opposite) are channeled downward instead of directly into the person they are speaking with as depicted in Figure 4. This is like how ceiling fans can displace accumulations of infectious particles during close-contact: for example, a mannequin placed directly underneath a ceiling fan was exposed to approximately 95% fewer aerosol particles from a simulated cough 5 feet away when the fan was on compared to when it was off [63]. Another question is how much of a ‘network effect’ occurs in a room if more people in the room use a wearable purifier or sleeping purifier (e.g. in berths), since a greater volume of air within the room will also get filtered.

**Figure 4:**
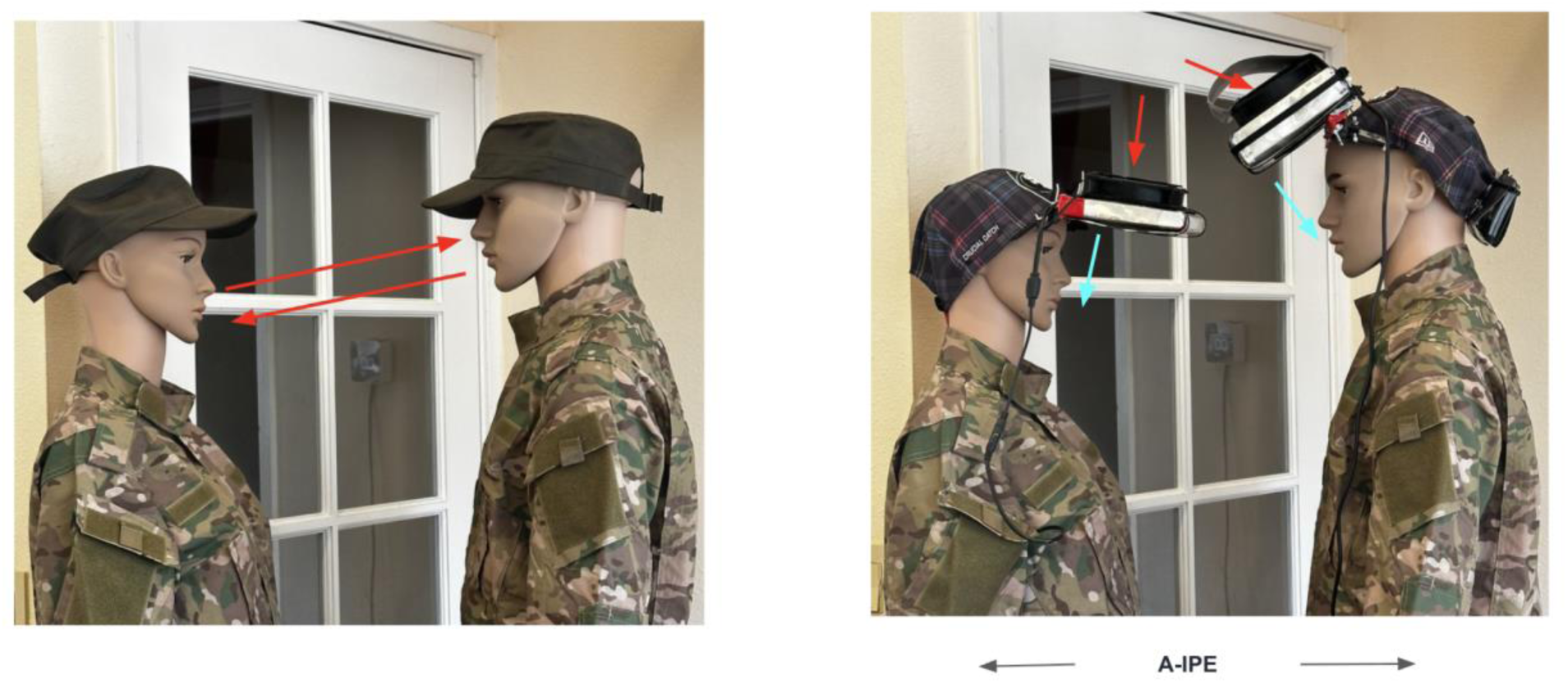
Depiction of “air jets” generated by exhalations from nose or mouth of another person and hypothesized yet unverified protection.

If a sufficiently airtight seal to the face can be formed using a traditional gas mask or N95, those may be more robust to windy conditions. However, such face seals often have leaks which are overcome by cap visor-mounted wearable air purifiers especially for indoor environments (when wind speed is limited). A transparent (half) face shield that extends from the eyes down to the mouth level can further prevent external aerosol mixing permitting eating and drinking. With the half face-shield, future prototypes could substantially reduce the size and hence weight of the filters. If a half face shield is included with the cap-visor mounted wearable air purifier device, one question is whether resilience to wind or exhalations from another person facing the wearer’s face can increases further.

Protection factor targets are approximate only, they rest on the studies cited, and complementary or contradictory data may alter them. Either more or less protection factor maybe needed in any specific situation.

## Conclusions

The handheld test method demonstrates a personally verifiable indoor protection factor of 5x-20x, 87%-98% for particulates at 0.3 μm at nose level of a mannequin using mass-manufacturable, facially unobstructed maskless protection devices (without burdens of masks).

In the US, the N95 mask incorporates manufacturing quality control administered by NIOSH, including spot checks of batches but not checks at the level of each individual mask. To transparently verify quality control for each individual device at time of assembly, a unique serial number can be assigned to each maskless device and a video recording of the test procedure performed on that device with a mannequin head can be retained for archived review by the end-user (wearer) independently of the manufacturer. Using the calibrated handheld particle counter, the protection provided to each mannequin (wearer) of the maskless device may be verified using the test protocol demonstrated on the mannequin in this study, like an optometrist verifies fit and function of eyeglasses in an eye shop, and adjustments made as necessary to ensure filtration efficiency, airspeed, and forward visibility of maskless devices.

Mask-free devices as tested were designed to permit access to the mannequin mouth for eating/drinking and all-day wearable protection from airborne infection and pollution indoors or outdoors but only without significant wind. As shown in Figure 5, although as-yet untested, the addition of an optional half- or full- face shield still permitting access to the mannequin mouth for eating/drinking is expected to improve protection during windy conditions outdoors by shielding from wind and guiding the filtered airflow towards the face.

**Figure 5:**
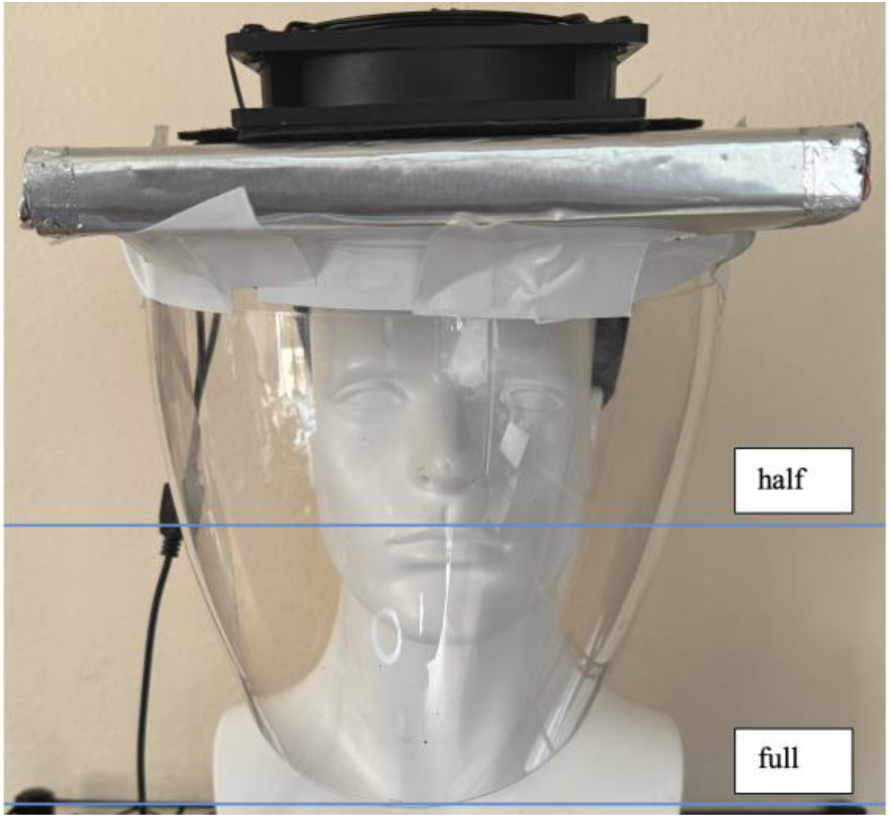
Adding a face shield, full (shown) or half to allow eating/drinking is expected to improve protection during outdoor windy conditions but as-yet untested

As debates continue about photochemistry induced pollution from UV for pathogen inactivation [64] and ionizers/electrostatic precipitation [65], the maskless devices described (based on fans and filtration) do not rely on those techniques.

Portable air purifiers can potentially provide 90–99% (10x-100x) protection factor from bioaerosols and airborne contagion with 10 ACH inside imperfectly sealed shelters/rooms if infiltration is 0.1-1.0 ACH (externally, internally, or uncontrolled entry/exit). Combined protection (multiplied) of maskless protection devices used with portable air purifiers of 99.98% (5000x) can in principle be achieved. To avoid restriction of movement (ROM) during uncontrolled contagion, submarines/ships/trains/hospitals may benefit from higher efficiency of both stackable air purifiers and sleeping purifiers doubling as air purifiers in berths with airborne contagion risk but limited floorspace.

## Supplementary Materials

### Air Filtration / Purifiers

The need for aerosol filtration to simultaneously address the dual concerns of toxic air pollution and airborne infection have been clear in California since at least 2020 when wildfires polluted the San Francisco Bay Area during the Covid pandemic [1]. In general, air pollution may arise from a variety of sources: accidents, wildfires, industrial sources, cooking, proximity to shooting guns (e.g. at a gun range), transportation (e.g. exhaust, brake dust). The Los Angeles fires that began on January 7, 2025 resulted in aerosol and vapor toxicity on a metropolitan scale with local residents experiencing measurable increases in pulmonary illness, myocardial infarction requiring emergency medical attention in the 90 day period following these fires [2]. These LA fires exposed the bottlenecks of masks necessitating indoor air cleaning for “collective protection.” Initially, on January 8, 2025, the California public health department (CDPH) posted on X for the public to use N95 masks to avoid inhalation of intense particulate pollution from the fires across the greater Los Angeles metropolitan area, as well to avoid respiratory viruses inside disaster shelters[3]. However, there was no public supply of N95 until a week later when city of Los Angeles announced N95s freely available at libraries on January 13 [4]. In the meantime, volunteers were handing out N95s and setting up air purifiers in disaster shelters (e.g. Pasadena Convention Center) yet many people could not wear these masks 24×7.

By Jan 15, in reaction to the LA fires, CDPH upgraded its indoor air cleaning recommendation [5] to meet or exceed the 2023 CDC recommendation of 5 air changes per hour in all occupied spaces [6]. These fires also triggered California’s statewide adoption of the 2023 CDC recommendation for 5 air changes per hour of aerosol filtration in all occupied spaces [7] which in principle provides a protection factor of approximately 10x from external pollution in a typical room assuming typical infiltration of 0.5 air changes per hour.

Not surprisingly, during the winter of 2025 air purifiers use in Delhi has grown representing the vast majority of air purifier sales in India according one electronics retailer [8], and the US embassy in Delhi replacing it’s the filters in its air purifiers [9]. However, as several observers noted the air purifiers installed at home do not permit protection during day-to-day activities outside the home [10].

### Individual Risk Model For Contagious Bioaerosols and Protection Factor

Individual infection risk (P) to susceptibles in a room or shelter with infectors is modeled according to the Wells-Reilly equation as [11]:

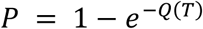

where Q(T) is the average number of infectious quanta inhaled by a susceptible individual during that time T. This formula may be interpreted as applying the Poisson distribution to the inhalation (arrival) of a random number of infectious quanta with average Q(T) into the respiratory system of the individual who is at risk. Q(T) can further broken down into the product:

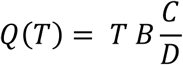

where B is the breathing rate of the individual at risk, C is the concentration of pathogen per unit volume in the room, and D is number of pathogens in one infectious quanta.

Under the well-mixed assumption, in steady state C can be deduced using the mass-balance relationship where the rate of exhalations of pathogen by infector(s) in the room (E) is exactly counterbalanced by the rate of removal of virus achieved by filtration or ventilation system in the room [equation 2 [12] and equation 3 of at equilibrium [13]]. The concentration of virus per unit volume (C) lowered by ventilation to satisfy the mass-balance condition to be function of (E) the total quanta emission rate of all infectors present and the rate of clean or filtered air flow introduced into the room (R) under the well-mixed assumption is:

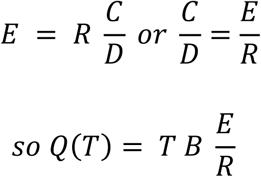

P can therefore be estimated from Q(T) as a function of E, T, and R – the quanta emission rate, duration of exposure, and ventilation/filtration rate which are independent of the number of susceptible people regardless of the size/area/volume of the room under the well-mixed assumption. The to the extent the well-mixed assumption is a valid approximation, same estimate applies for a shelter, a ship, a submarine, or an airplane regardless of its size. IPE reduces the exposure by the IPE’s fixed protection factor (F) e.g. 20x for N95, 100x for N99, etc. The effect of IPE can be modeled as reducing Q(T) by a factor of F in the denominator, equivalent to scaling the ventilation rate:

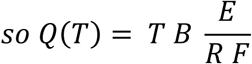

Figure S1 and Figure S2 show F as a function of E (quanta emission rate), R (ventilation or filtration rate) for T = 1 hour (e.g. classroom or lunch with infector) and T = 8 hours (e.g. sleeping in same room as infectors). R ranges from 1 to 10,000 CFM (assuming breathing rate B = 0.52 cubic meters per hour. Quanta emission rate ranges from 0 to 1000 representative of values found in [13] and [14]. Both Figure S1 and Figure S2 illustrate the rise in infection risk from 0-100% as a function of infector’s quanta emission rate and countervailing drop in risk as clean air airflow rate rises. The rapid increase of infection risk when people are together for prolonged periods such a 8 hours (Figure S3), as opposed to just one hour (Figure S2), can be seen clearly from the dependence of individual risk on total airflow rate where even a thousand CFM maybe sufficient to keep the infection risk low for 1 hour, much more exceeding 5,000 CFM maybe needed for 8 hours depending on the projected quanta emission rate under the well-mixed assumption.

**Figure S1:**
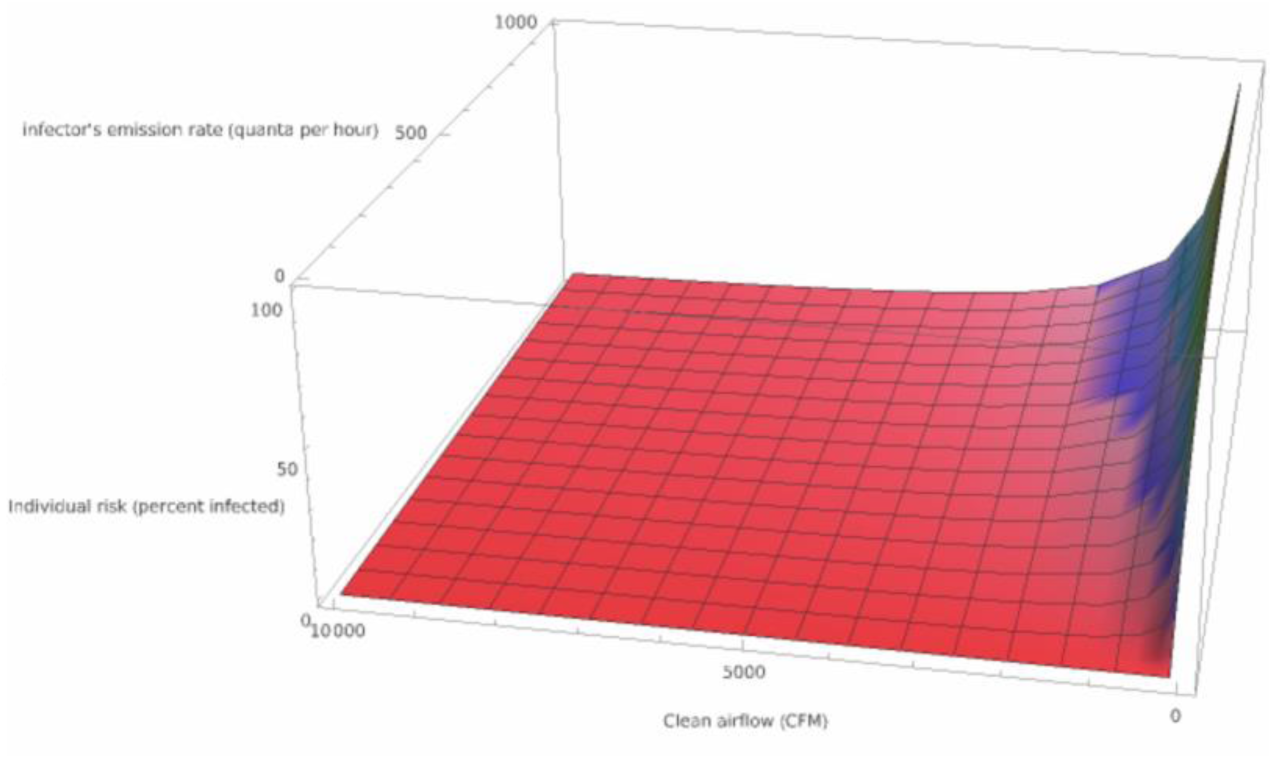
Individual infection risk over 1 hour versus infector quanta emission rate (per hour) and filtration/ventilation airflow rate (cubic feet per minute)

**Figure S2:**
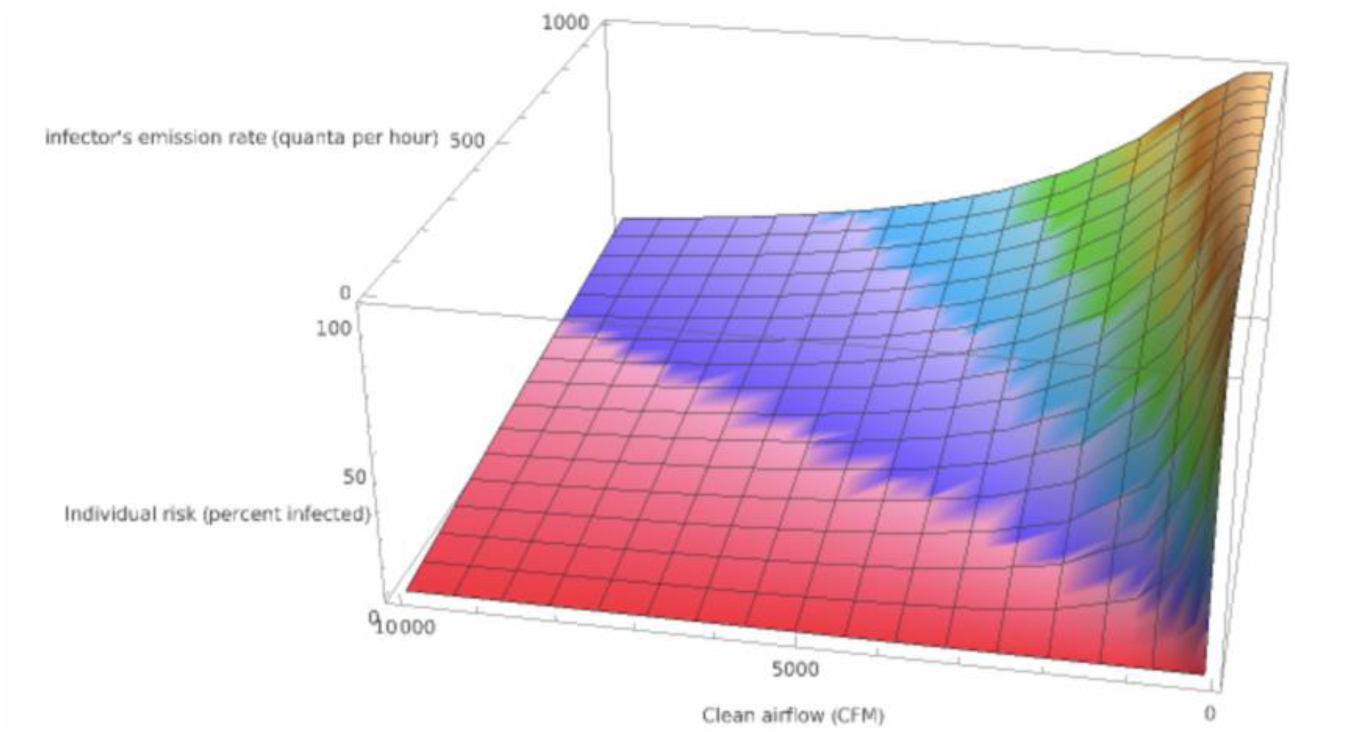
Individual infection risk over 8 hours versus infector quanta emission rate (per hour) and filtration/ventilation airflow rate (cubic feet per minute)

The effect of adding individual/personal protective equipment (IPE/PPE) either to the susceptible(s) or to the infector(s) may therefore be visualized on Figure S1 and Figure S2 by simply “boosting” the ventilation (R) by the IPE’s protection factor F. By combining the ventilation (R) with an IPE protection factor F=20 (95%), the 8-hour risk rapidly drops as shown in Figure S3, to levels below the 1-hour exposure of Figure S4 since time (T) is in the numerator of the expression for Q(T).

**Figure S3:**
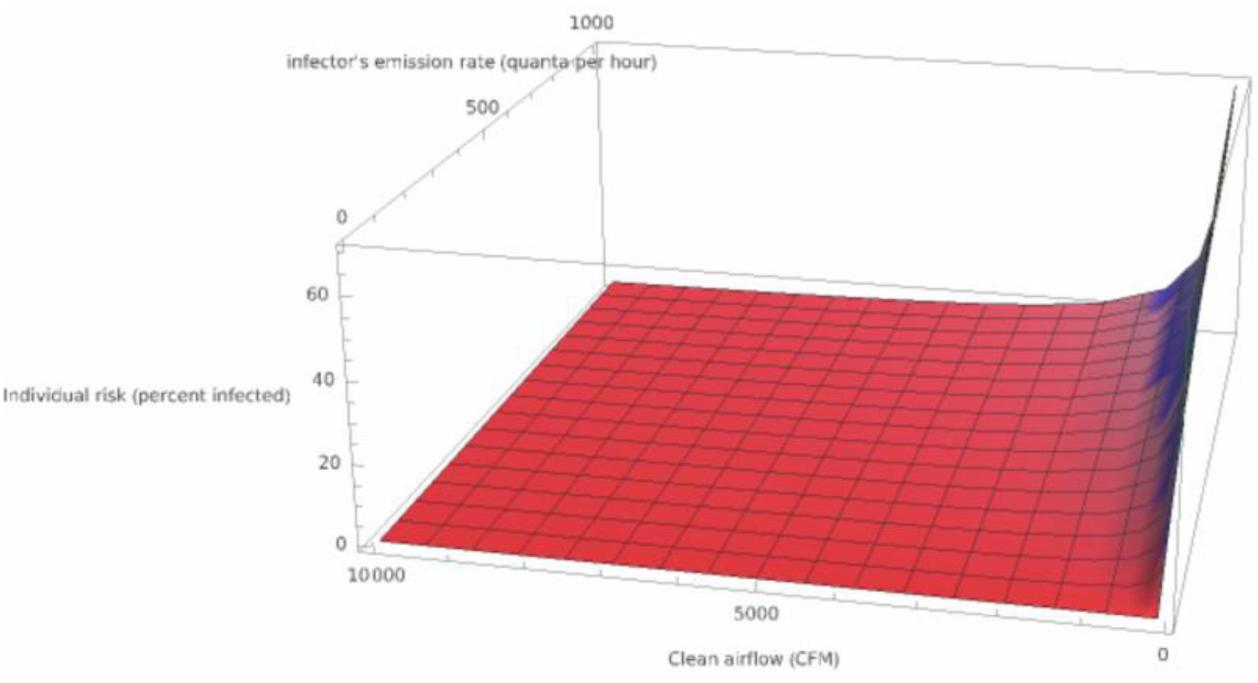
Individual infection risk over 8 hours versus infector quanta emission rate (per hour) and filtration/ventilation airflow rate (cubic feet per minute) with added protection factor F=20 (95%)

The well-mixed assumption may not always apply even with good ventilation when people work or sit with others in close proximity (e.g., while eating lunch less than 3 feet from each other for 30 minutes) especially in the “jet zone” from an infector [15] or who have fleeting contact in hallways are at risk for pathogen transmission [16]. At such short distances, the infector’s emissions do not have enough time to dilute inside the room before reaching a susceptible person nearby, particularly during prolonged exposure (e.g., consider someone smoking a cigarette). Empirical evidence supports this distance-dependent risk: a study of COVID-19 transmission in prisons found that inmates were much more likely to be infected if they were exposed to another infected person within their own cell than within the same cell block, possibly because exposure within the same cell is much closer and for a longer duration [17]. Also a study of digital contact tracing of SARS-CoV-2 among 7 million close contacts (e.g., within about 6 feet) similarly concluded that although many exposures were short in duration, recorded transmissions typically resulted from exposures lasting from an hour to multiple days [18]. In summary, infection risk is both bioaerosol concentration-dependent and exposure duration-dependent, which to reduce transmission risk in scenarios with infection risk, and needs a high degree of continuous collective and personal protection suitable for extended-use [19].

### Design considerations

Each of the maskless devices is based on an air processing assembly that at minimum consists of filter(s), a fan, a fan grille, and optionally an aluminum case/fan-guard to shield the fan from rain or other elements. The filter case was chosen to be cut from aluminum sheet metal to minimize temperature-dependent off-gassing that can occur in alternative materials such as cardboard or plastic. In devices needing a counterweight, it was also made from aluminum. Each of the elements were attached using duct tape or cable ties. “Pro” and “Max” included L-brackets to raise the air processing assembly above the brim of the hat. All devices used friction hinges attached to the brim of the hat to finely tune the angle of the airflow relative to the vertical while maintaining forward visibility.

The direction of filter pleats was perpendicular to the direction facing the wearer of the device. Although HEPA filters could be used, they would have required two or more of the most powerful case fans available stacked in series to generate the static pressure needed to overcome the higher level of air resistance, making them heavier, larger, and less wearable. Use of filters cut from N95 masks did not result in a wearable maskless device because of their air resistance [20]. This reflects the competing physical challenges for a wearable, maskless (unsealed) device to achieve stringent targets of 95% filtration at 0.3 µm at nose level:

- Filtration-wearability vicious cycle: each extra particulate contamination filter that is added in an airstream (in series) to increase filtration efficiency also introduces air resistance, which in turn decreases airflow velocity, and therefore must be then counteracted by increasing fan airflow and air pressure, which in turn increases noise generated, size, weight, and power consumed making the device unwearable (the ‘filtration-wearability challenge’).
- Intermixing of submicron particles: When the face is unobstructed, there is a relatively large distance (approximately 2-3 inches or more) between the filter (positioned above or below the breathing zone) and the nose that allows for intermixing of the filtered air with surrounding contaminated air as it travels, by the time it reaches the nose level. This intermixing occurs near the boundaries of the filtered airflow in any unsealed gaps possibly through a variety of aerodynamic and fluidic processes such as entrainment, turbulence, diffusion, pressure differential, shearing, etc. potentially in a particle-size dependent manner. The intermixing effects particles having a size of about 0.3 μm since a moving airstream also allows particles to enter (mix) from adjacent, stationary contaminated air in unsealed systems with large gaps.

During the construction, it was observed that in some cases MERV 13-14 filters may have spatially non-uniform filtration efficiencies for particles sized 0.3 µm within the 6”x9” cuts. In other words, the filtration efficiency may be much lower in some locations than others even within the same filter. In such cases either replacement or unit testing of different locations on the original 20”x20” filters was necessary to select 6”x9” sections that did not have dips in filtration efficiency, to avoid non-uniform or asymmetric filtration efficiency across different locations on the face especially in “Pro” and “Max” with two stacked MERV filters.

### Handheld Test Procedure for Air Changes Per Hour (ACH)

This test procedure to measure ACH is similar to that described Supplemental Appendix of [19]. The air filtration in air changes per an hour (ACH) was measured for a test aerosol (smoke generated by burning marshmallow) in a test room of volume 2,961 cubic feet with windows and doors closed and central ventilation system turned off. In contrast to industry-standard test of air cleaners tested in rooms of about 1000 cubic feet where mixing effects maybe distorted by small volume [21], the use of a test room of about 3,000 cubic feet permits observation of the mixing effects from higher airspeed at the fan inlet or outlet. Test aerosol/vapor concentrations were measured using OPC1. The air changes per hour (ACH) for with test aerosol is determined by fitting the shape of the decay curve and asymptote of measurements from the OPC1 and PID in the test room using on the following equation based on the model in [13].

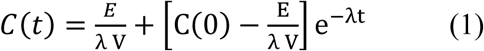

where:

C(t) = concentration of aerosol at time t
C(0) = initial concentration of aerosol at time 0
E = rate of aerosol infiltrating into the room (per hour) e.g. sourced from outside or inside the room or from desorption inside the room
V = volume of the room
λ = Air changes per hour (ACH) of the air cleaning (filtration)
e = base of the natural logarithm

The aerosol concentration measurements representing C(t) were fitted by a computer program written in Python to obtain the best approximation for ACH (λ) and the rate of indoor leak (E/V) by minimizing the sum of percent error across each measurement. The volume (V) was measured to be 2,961 cubic feet, from which the clean air delivery rate (CADR in units of cubic feet per minute) for the test aerosol was estimated by the following formula:

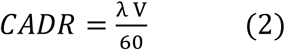

